# Early immune cell development precedes gastrulation in annual killifish

**DOI:** 10.64898/2026.06.25.734555

**Authors:** Sydney M. Sattler, Nicholas C. Lammers, Sebastian V. Hriscu, Clare L.T. Booth, Cole Trapnell, Philip B. Abitua

## Abstract

During embryogenesis, cell types arise in a predictable order because developmental regulators act sequentially. But how evolutionary changes in morphogenesis reshape the signaling environments that activate these regulators remains unknown. Across vertebrates, primitive myeloid cells emerge from bone morphogenetic protein (BMP)-patterned ventral mesoderm through a conserved regulatory program. Here we show that in the annual killifish, *Nothobranchius furzeri*, a vertebrate with highly derived embryogenesis, neutrophils emerge prior to gastrulation, before ventral mesoderm has formed. Vascular progenitors arise later from ventral mesoderm, whereas myeloid progenitors are largely absent from this tissue. BMP inhibition abolishes pre-gastrula neutrophil specification, while disruption of Nodal-dependent mesendoderm formation does not. These findings reveal that conserved cell type programs can be redeployed within an altered embryonic architecture.

## Main Text

Cell fate specification depends not only on the transcription factors a cell expresses, but also on the developmental context in which that cell resides. In vertebrate embryos, lineage-specifying gene regulatory networks (GRNs) are activated within a stereotyped sequence of morphogenetic events, including axis formation, gastrulation, and organogenesis, which establish tissue organization and signaling environments that constrain when and where cell types can emerge (*1*). These constraints could reflect context dependence inherent to the GRN. Alternatively, they could arise from evolutionarily contingent features of embryonic architecture that position the relevant signals within particular tissues. In the latter case, where and when a cell type emerges would be determined by how the embryo is built rather than by the network logic itself.

Annual killifish embryos provide a natural system for testing whether GRN deployment is necessarily coupled to tissue-level morphogenesis, as their early embryogenesis diverges substantially from that of other vertebrates. Native to ephemeral pools in southeastern Africa that evaporate each dry season, the annual killifish, *Nothobranchius furzeri* (*N. furzeri*), evolved under intense selective pressure (*2*), producing a highly derived mode of early development (*3*, *4*). Following cleavage, deep blastomeres disperse as individual cells across the yolk surface and remain motile; the embryo lacks discernible tissue organization (Fig. 1A) (*3–5*). Despite this, β-catenin and Nodal initiate mesoderm formation within a small cluster of cells known as the incipient aggregate (*6*). Notably, mesodermal markers appear before ectodermal markers, inverting the canonical germ layer sequence observed in other vertebrates (*6*).

**Fig. 1.**
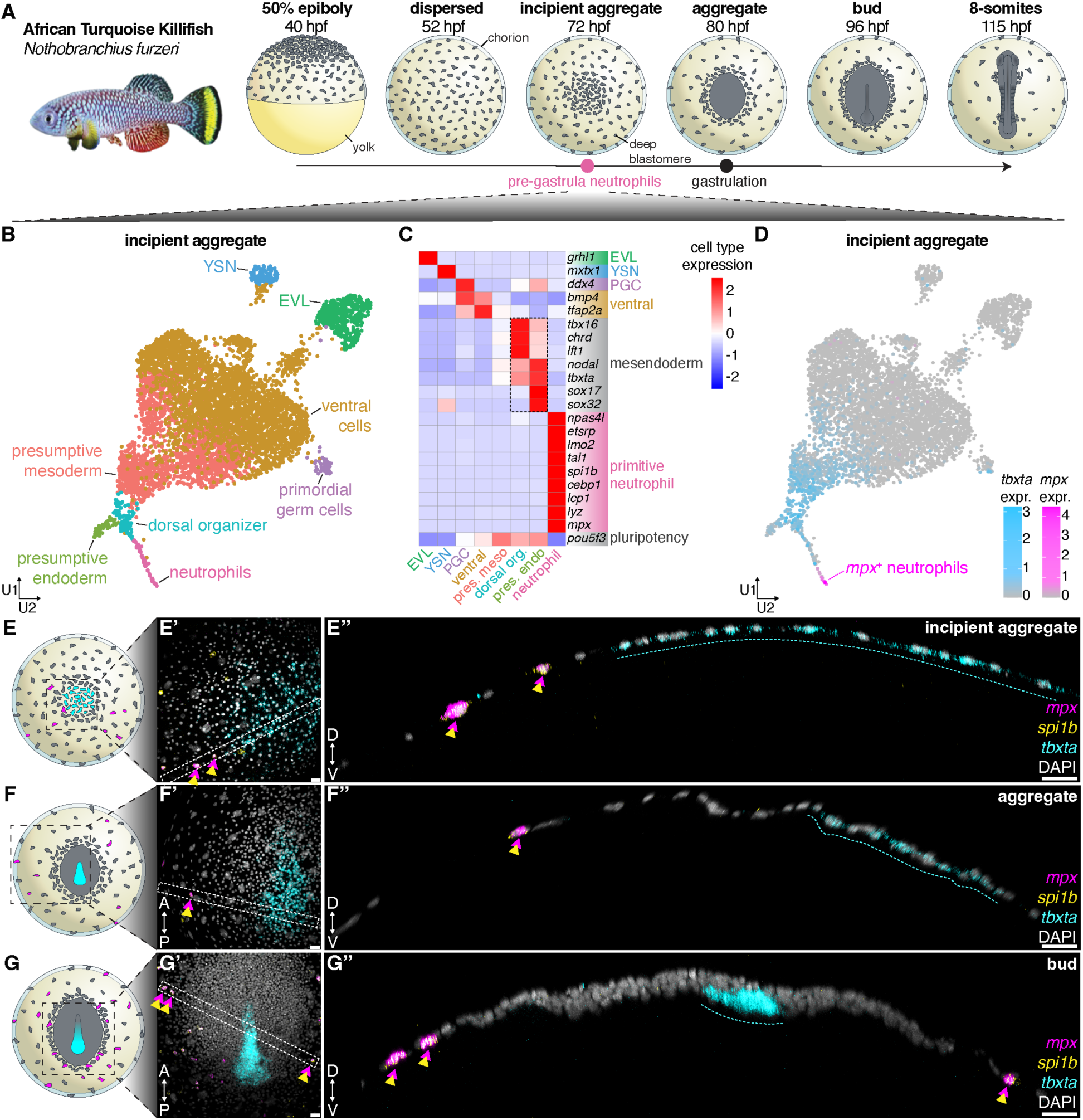
Neutrophil GRN deploys before gastrulation. (**A**) Schematic of early *N. furzeri* embryogenesis. (**B**) UMAP of incipient aggregate cell types. (**C**) Heatmap of gene-scaled average expression of selected marker genes across cell types in (B). (**D**) UMAP expression of *tbxta* (cyan) marking presumptive mesoderm and *mpx* (magenta) marking neutrophils. (**E** to **G**) Stage schematics show inset regions for (E’) to (G’). (**E’** to **G’’**) *In situ* HCR shows the expression patterns of presumptive mesoderm (*tbxta*, cyan), myeloid progenitors (*spi1b*, yellow), and neutrophils (*mpx*, magenta) at the incipient aggregate, aggregate, and bud stages. White dashed rectangles in (E’) to (G’) indicate positions of orthogonal ZX slices shown in (E”) to (G”). Double-headed magenta and yellow arrowheads mark neutrophils coexpressing *spi1b* and *mpx*; dashed cyan lines mark *tbxta^+^*presumptive mesoderm and its derivatives. EVL, enveloping layer cell; YSN, yolk syncytial nuclei; PGC, primordial germ cell; pres. meso., presumptive mesoderm; dorsal org., dorsal organizer; pres. endo., presumptive endoderm; hpf, hours post-fertilization. Scale bars: 30 µm.

This prompted us to ask whether other cell types might also emerge outside their typical developmental order. By profiling early embryogenesis in *N. furzeri* and comparing myeloid development with zebrafish, we found that primitive myelopoiesis, the earliest wave of innate immune cell specification, occurs unexpectedly early. In other vertebrate models, primitive myelopoiesis is initiated during segmentation within BMP-patterned ventral mesoderm, giving rise to embryonic macrophages and neutrophils (*7*, *8*) through a conserved developmental program involving *npas4l*, *etsrp/etv2*, and *spi1* family regulators (*9–15*). In contrast, neutrophils differentiate prior to gastrulation in *N. furzeri*, revealing that primitive myelopoiesis is not intrinsically coupled to ventral mesoderm.

## Primitive myeloid GRN deployment precedes gastrulation in *N. furzeri*

To characterize cell fate specification during early embryogenesis, we performed single-cell RNA sequencing (scRNA-seq) on *N. furzeri* embryos at the incipient aggregate stage, prior to gastrulation. In addition to the previously characterized dorsal organizer and presumptive mesodermal populations (*6*), we identified transcriptionally distinct primordial germ cells (PGCs), ventral embryonic cells, presumptive endoderm, and extraembryonic enveloping layer (EVL) cells, as well as yolk syncytial nuclei (YSN) (Fig. 1B and table S1).

Although tissue organization is largely absent at this stage (*6*), the dorsal-ventral (D-V) axis has already been established, with dorsal and ventral signaling programs expressed in transcriptionally distinct cell populations (Fig. 1C and fig. S1A). Germ layer segregation, however, remains incomplete, as cells expressing endodermal markers (*sox32, sox17*) retain expression of mesodermal genes (*chrd, lft1, tbxta*) (Fig. 1C and fig. S1B). Additionally, neuroectodermal markers such as *otx2* are absent (fig. S1C). The pluripotency factor *pou5f3* (*16*) remains broadly expressed across mesendodermal populations but is downregulated in extraembryonic and PGC clusters (Fig. 1C and fig. S1D). These data establish that axial patterning has initiated at the incipient aggregate stage, but the germ layers are not yet transcriptionally distinct.

Despite the absence of definitive germ layers, we identified a distinct cluster of cells expressing the mature neutrophil marker *mpx* (Fig. 1, C and D, and table S1) (*17*). This cluster expressed key components of the primitive neutrophil GRN, including *npas4l*, *etsrp*, *spi1b*, and *cebp1* (Fig. 1C and table S1), had downregulated *pou5f3*, and expressed additional differentiated neutrophil markers including *lcp1* and *lyz*, consistent with mature neutrophil identity (Fig. 1C and table S1) (*8*, *10*, *18*, *19*). Collectively, this transcriptional profile identifies differentiated neutrophils at the incipient aggregate stage, well before the typical onset of primitive neutrophil-specifying factors during segmentation in zebrafish and other vertebrates (*8*, *9*).

We next examined the spatial distribution of these cells during aggregate formation. At the incipient aggregate stage, deep blastomeres begin to coalesce into a monolayer, with *tbxta*-expressing mesendodermal cells occupying the center of the incipient aggregate (Fig. 1, E to E”). Concurrently, neutrophils coexpressing *spi1b* and *mpx* are already present at the periphery of the forming *tbxta*^+^ incipient aggregate and on the yolk surface where cells remain dispersed (Fig. 1, E to E”). As development proceeds, *tbxta*^+^ mesoderm consolidates further (Fig. 1, F to F”), and by the bud stage, mesodermal cells have ingressed and posterior axis elongation has begun (Fig. 1, G to G”). Thus, neutrophils emerge before gastrulation has generated organized germ layers, including the ventral mesodermal tissue from which primitive myeloid cells normally arise (*8*, *9*, *15*).

## Neutrophils emerge sparsely and asynchronously at the aggregate periphery

To visualize neutrophils *in vivo*, we generated a CRISPR knock-in of eGFP to the endogenous *lcp1* locus, marking mature myeloid cells (*7*, *8*), and combined this reporter with a ubiquitously expressed nuclear marker to provide spatial context for *in toto* imaging. Live light-sheet microscopy from the start of epiboly through aggregation confirmed that a subset of cells activated *lcp1* before gastrulation and enabled direct lineage tracing of neutrophil emergence (movie S1).

*Lcp1*^+^ neutrophils were typically first detected as sister cell pairs, with reporter activation occurring within minutes of division (Fig. 2A). The timing of sister cell pair appearance spanned a broad 12-hour window beginning at incipient aggregate formation, indicating asynchronous differentiation (Fig. 2A). Although an earlier common progenitor cannot be excluded in cases where overlapping cells could no longer be unambiguously distinguished, tracing progenitors prior to reporter activation revealed no common ancestor within the imaging window (Fig. 2A). These observations suggest that pre-gastrula neutrophils do not arise from a common fate-restricted lineage but instead emerge independently from dispersed deep cells.

**Fig. 2.**
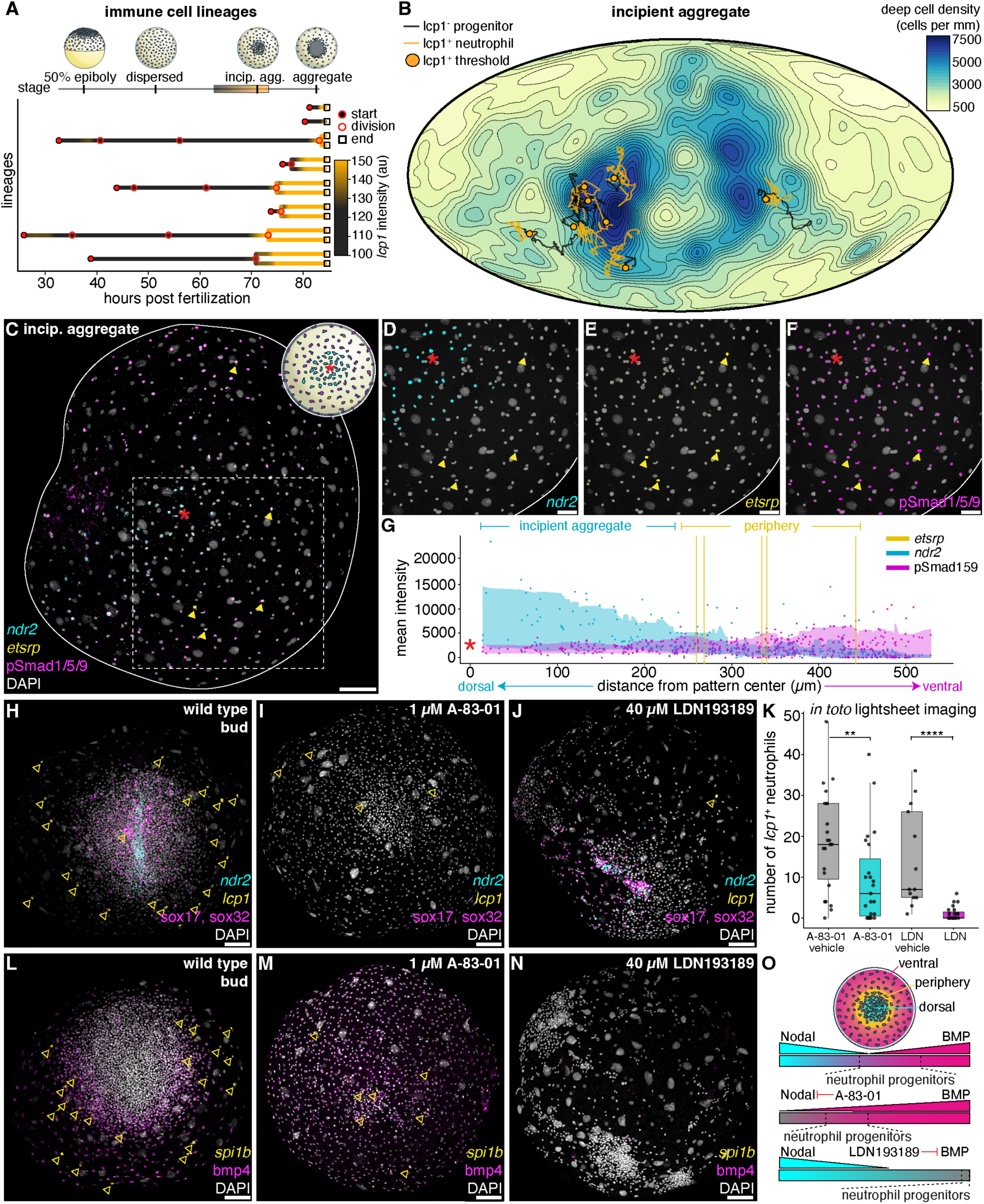
Peripheral neutrophil specification requires BMP. (**A**) Neutrophil lineages colored by *lcp1* intensity (closed circles, lineage start; open circles, cellular divisions; open squares, lineage end). Embryo schematics depict developmental stages. The grey to orange rectangle marks the developmental window analyzed in (B). (**B**) Mercator projection from 64-74 hpf colored by deep cell density (nuclei/µm^2^) with neutrophil lineages overlaid. Tracks show *lcp1*^-^ progenitors (dark gray) transitioning to *lcp1*^+^ neutrophils (orange); orange circles mark reporter activation. (**C**) Embryo schematic and *in situ* HCR+IHC at the start of the incipient aggregate stage (66 hpf) marking the dorsal organizer (*ndr2*, cyan), BMP-active ventral cells (pSmad1/5/9, magenta) and myeloid progenitors (*etsrp*, yellow). Red asterisk marks the dorsal patterning center; dashed box indicates the region quantified in (D) to (F). (**D** to **F**) Nuclear masks pseudocolored by quantified *ndr2*, *etsrp*, or pSmad1/5/9 intensity. (**G**) Radial profiles of *ndr2*, *etsrp*, and pSmad1/5/9 intensity relative to the dorsal patterning center; yellow lines mark *etsrp*^+^ cells. (**H** to **J**) *In situ* HCR marks mesoderm (*ndr2*, cyan), endoderm (*sox17, sox32*, magenta) and neutrophils (*lcp1*, yellow) in embryos treated from 48-72 hpf with DMSO/HCL control (n = 15/17) (**H**), 1 µM Nodal inhibitor A-83-01 (n = 20/20) (**I**), or 40 µM BMP inhibitor LDN-193189 (n = 17/17) (**J**), and fixed at 96 hpf. (**K**) Quantification of *lcp1*^+^ neutrophils from *in toto* light-sheet imaging of *NLS-mScarlet*; *eGFP*-*lcp1* embryos treated as in (H) to (J). (**P = 9e^-3^; ****P = 1e^-5^; n = 82). (**L** to **N**) *In situ* HCR marks neutrophils (*spi1b*, yellow), myeloid progenitors (*etsrp*, cyan), and ventral mesoderm (*bmp4*, magenta) in embryos treated as in (H) to (J) (n = L:20/20, M:14/17, N:16/16). (**O**) Model of early neutrophil progenitor positioning relative to Nodal and BMP signaling. Incip. agg., incipient aggregate; au, arbitrary units. Scale bars: 100 µm (C), (H) to (J), (L) to (N), 50 µm (D) to (F).

During the incipient aggregate stage, a 2D Mercator projection of the embryo surface showed that deep cell density was highest within the forming aggregate (Fig. 2B and movie S2). *Lcp1*^+^ neutrophils emerged at the periphery of this high-density domain, rather than within its center or across the yolk surface (Fig. 2B, fig. S2C and movie S2). Neutrophils did not arise from a contiguous field; instead, they formed in a salt-and-pepper pattern along the periphery of the aggregate (Fig. 2B and movie S2). Together, these findings demonstrate that pre-gastrula neutrophils differentiate asynchronously within a spatially restricted domain at the aggregate periphery.

## BMP signaling defines a permissive zone for neutrophil specification

Across vertebrates, opposing gradients of BMP and Nodal direct both embryonic axis formation and subsequent primitive myeloid specification (*11*, *13*, *15*, *20–22*). In *N. furzeri*, at the onset of incipient aggregate formation, the Nodal ligand *ndr2* marked the dorsal organizer (*6*), whereas BMP activity, detected by pSmad1/5/9 staining, broadly labeled the surrounding dispersed deep cells, together establishing the D-V axis (Fig. 2, C, D, F, and G). Early neutrophil progenitors, identified by expression of the upstream regulator *etsrp* (*13*), were enriched at the incipient aggregate periphery within a region of active BMP signaling and outside the *ndr2*^+^ organizer domain (Fig. 2, C, E, and G). These observations identify a BMP-active peripheral domain surrounding the forming organizer as the site of early neutrophil specification.

To test the functional requirements of these pathways, we inhibited Nodal signaling during the window of early neutrophil specification. Treatment with A-83-01, a potent inhibitor of Activin/Nodal receptors (*23*), blocked aggregation and mesendoderm specification (*6*), as evidenced by the loss of *ndr2*, *sox17*, and *sox32* expression (Fig. 2, H and I). Nodal-inhibited embryos were severely ventralized, with ectopic *bmp4* expression throughout the blastoderm, consistent with loss of organizer function (Fig. 2, L and M). Despite these defects, reduced numbers of *lcp1*^+^ neutrophils still emerged (Fig. 2, H, I, K to M), indicating that Nodal-dependent mesendoderm formation is not required for pre-gastrula neutrophil fate specification.

In contrast, inhibition of BMP signaling using LDN-193189 (*24*) resulted in severe dorsalization marked by the complete loss of *bmp4* expression (Fig. 2, L and N). Mesendodermal cells (*ndr2*^+^, *sox17*^+^, and *sox32*^+^) failed to resolve into a coherent axial domain, remaining diffuse and disorganized (Fig. 2, H and J). Under these conditions, *spi1b*^+^, *lcp1*^+^ coexpressing neutrophils were strongly reduced or eliminated (Fig. 2, H, J to L, and N), demonstrating that BMP signaling is required for early neutrophil specification. These findings indicate that early neutrophil specification occurs within a BMP-dependent permissive zone at the aggregate periphery and can occur without Nodal-mediated mesendoderm formation (Fig. 2O).

## Neutrophils specified prior to gastrulation exhibit functional immune behavior

We next asked whether early neutrophils are functionally competent. At the bud stage, *lcp1*^+^ neutrophils patrolled across the yolk at speeds approximately two-fold greater than *lcp1*^-^ deep cells (fig. S2, A, B, and D, and movie S3), consistent with the rapid motility of vertebrate neutrophils (*25–27*). They also displayed dynamic protrusive activity and amoeboid morphology characteristic of vertebrate neutrophils, neither of which was observed in *lcp1*^-^deep cells (Fig. 3A, fig. S2, A, B, and D, and movie S3) (*25–27*). To determine whether early neutrophils can mount an immune response, we inflicted a stab wound at the bud stage and monitored cell behavior by live light-sheet imaging. *Lcp1*^+^ neutrophils migrated toward the wound, dwelled at the injury site, and later exited by reverse migration (Fig. 3, B to C’, and movie S4), consistent with neutrophil recruitment and resolution behaviors described in zebrafish (*28*). *In situ* hybridization chain reaction (HCR) confirmed that *lcp1*^+^, *mpx*^+^ coexpressing neutrophils accumulated at the wound site (Fig. 3D and fig. S3). These results demonstrate that early *N. furzeri* neutrophils can mount a robust inflammatory response to tissue injury. To our knowledge, this represents the earliest immune response reported in a vertebrate embryo (*7*, *9*, *29*, *30*).

**Fig. 3.**
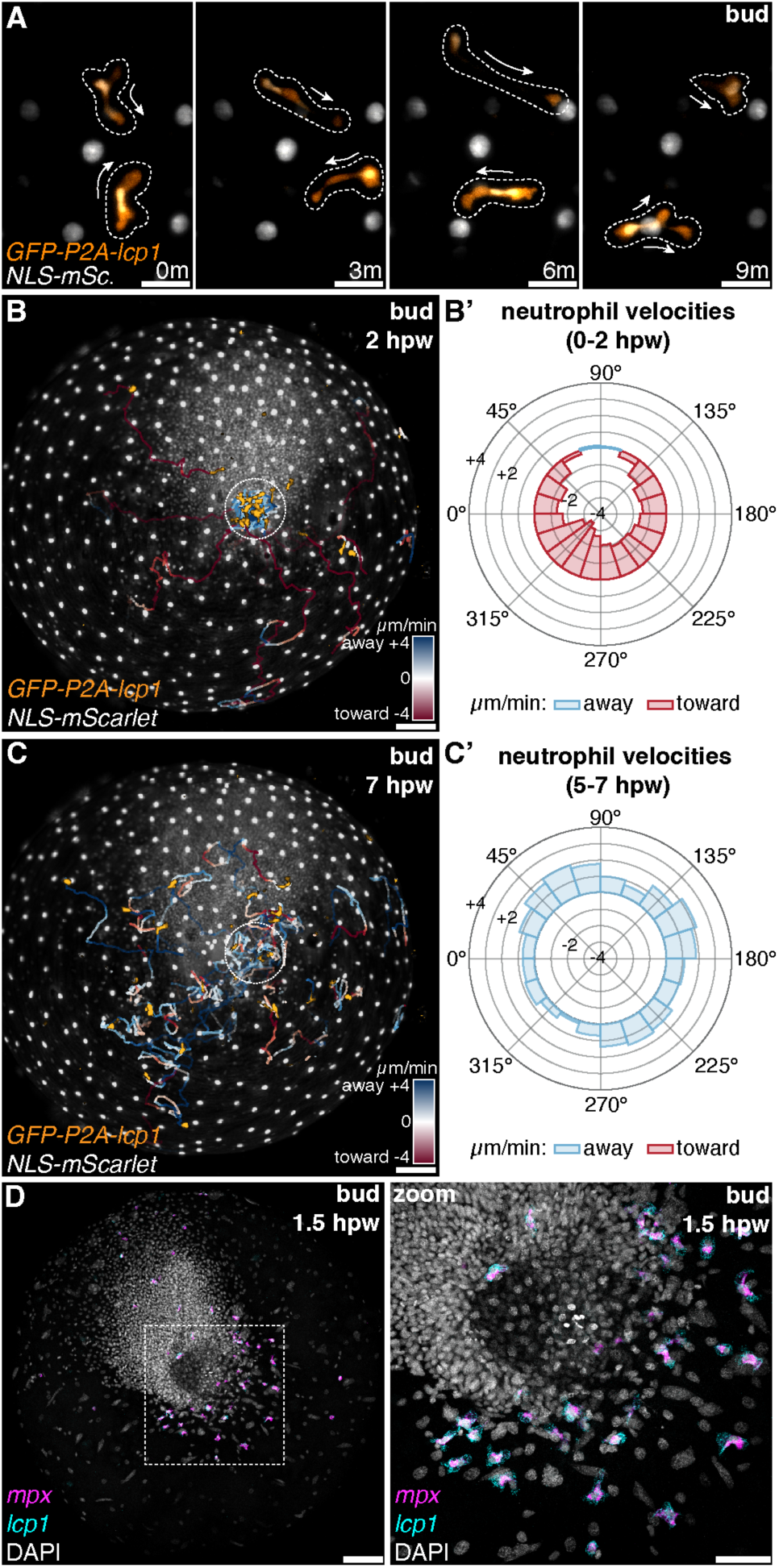
Neutrophils function before organogenesis. (**A**) Time-lapse images showing the morphology and behavior of two neutrophils (*eGFP*-*lcp1*, orange; pan-nuclei, white) over 9 minutes during the bud stage. Dashed lines outline cells. White arrows indicate migration direction. (**B** and **C**) Tracks of *lcp1+* neutrophils from 1.5-2 hpw (B) and 6.5-7 hpw (C), colored by movement toward (red) and away (blue) from the wound. Dashed circle marks stab wound location. (**B’** and **C’**) Rose plots showing neutrophil velocities (µm/min) toward (red) and away from (blue) the wound between 0-2 hpw (B’) and 5-7 hpw (C’). (**D**) *In situ* HCR marks neutrophils (*mpx*, magenta; *lcp1*, cyan) localized to the wound site 1.5 hours after bud-stage injury (n = 13/15). Dashed box indicates zoomed-in region. hpw, hours post-wounding. Scale bars: 25 µm (A), 50 µm, zoom of (D), 100 µm (B) to (D).

## Primitive myelopoiesis is uncoupled from the anterior lateral plate mesoderm in *N. furzeri*

In zebrafish, primitive myelopoiesis originates in the anterior lateral plate mesoderm (ALPM), where *spi1b*^+^ myeloid progenitors arise adjacent to vascular lineages under shared BMP signaling and give rise to macrophages followed by neutrophils (*7*, *8*, *11*, *13*, *15*, *18*, *22*, *31*). Having shown that functional neutrophils emerge prior to gastrulation in *N. furzeri*, we next asked whether the ALPM contributes to primitive myelopoiesis as it does in zebrafish.

To explore this question, we generated additional scRNA-seq datasets from the dispersed, bud, and 8-somite stages and integrated them with the incipient aggregate dataset, producing the first atlas of early killifish embryogenesis, available through SKiRA (Single-cell Killifish RNA Atlas) (fig. S4 and table S2 to S5). Using marker gene expression, we identified lateral plate mesoderm (LPM) populations and their derivatives (fig. S5 and table S6). LPM-derived cell types were not detected at the incipient aggregate or bud stages, despite the presence of the pre-gastrula neutrophil lineage (fig. S5, F and G). By the 8-somite stage, LPM derivatives were present, including *kdrl*^+^ vascular progenitors, *gata1a*^+^ erythroid progenitors, and *irf8*^+^ macrophages (fig. S5H and table S7) (*32–36*). At this stage, *spi1b* expression was restricted to mature neutrophil and macrophage clusters, and no discrete *spi1b*^+^ myeloid progenitor population was identified (fig. S5I). These observations suggest that *N. furzeri* lacks a *spi1b*^+^ ALPM-associated myeloid progenitor domain equivalent to that in zebrafish.

To compare the spatial distribution of these cell populations in zebrafish and killifish, we performed *in situ* HCR for markers of the ALPM (*nkx2.7*), vascular progenitors (*etsrp*), and myeloid cells (*spi1b*) (*13*, *15*, *37*). In both species, the *nkx2.7*^+^ ALPM first appeared as a horseshoe-shaped territory at the bud stage and narrowed into bilateral stripes during segmentation (Fig. 4, A to D, and fig. S6) (*37*). In zebrafish, *etsrp*^+^ vascular progenitors resolved into a contiguous domain, while *spi1b*^+^ myeloid progenitors formed immediately lateral to this domain by the 2-3 somite stage (Fig. 4C). As previously reported (*8*, *15*, *38*, *39*), *spi1b*+ myeloid progenitors then converged toward the midline (fig. S6A) before migrating onto the yolk (fig. S6C). In killifish, *etsrp*^+^ vascular progenitors occupied a similar position within the ALPM but formed a looser network of cells (Fig. 4D). In contrast to zebrafish, no ALPM-associated *spi1b*^+^ myeloid progenitor domain was observed at any stage examined; instead, spi1b+ cells were restricted to the yolk, consistent with the pre-gastrula-derived myeloid population (Fig. 4, B, D, H, and fig. S6B).

**Fig. 4.**
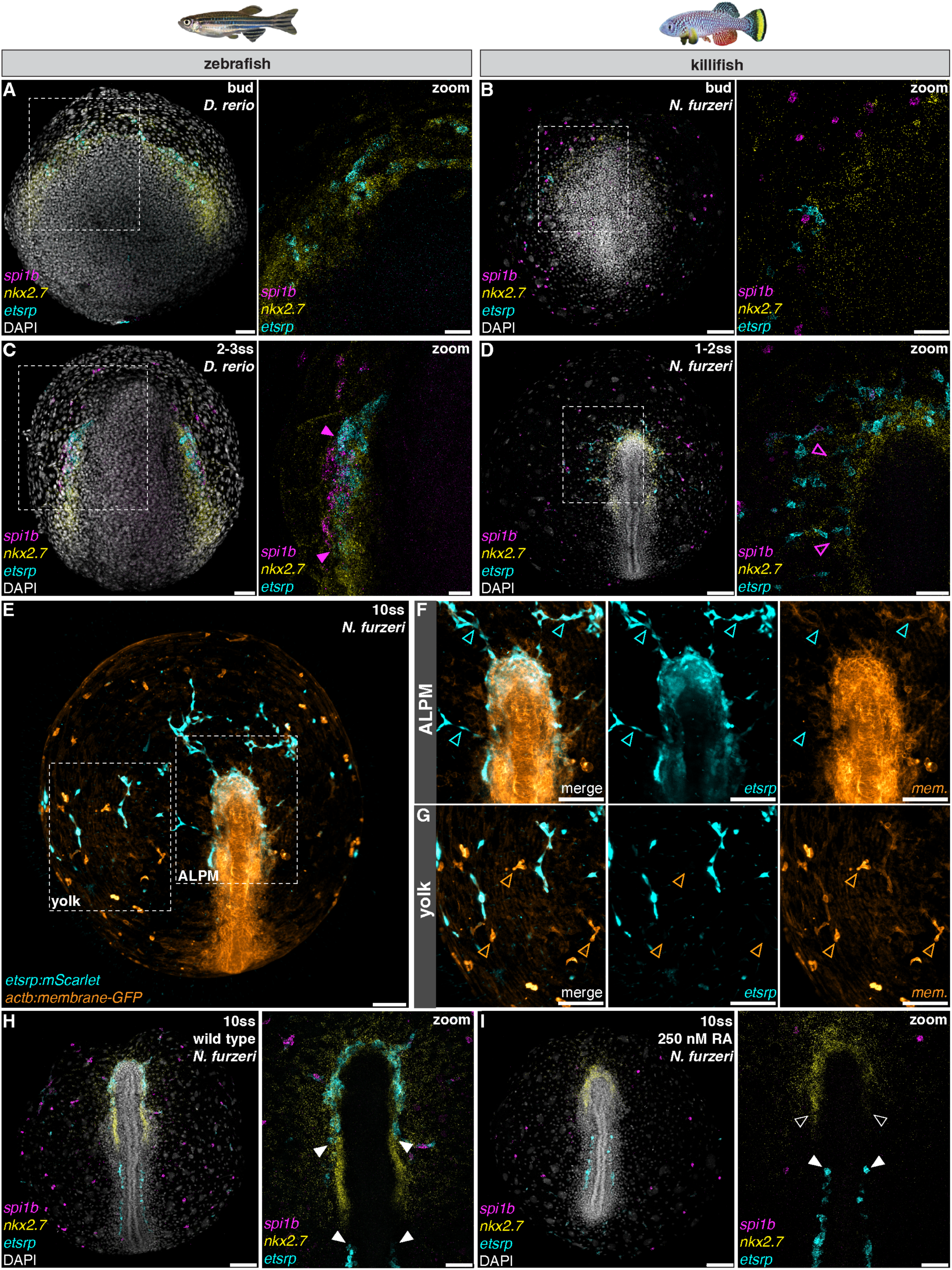
ALPM-independent myelopoiesis in killifish. (**A** to **D**) *In situ* HCR marks the ALPM (*nkx2.7*, yellow), vascular progenitors (*etsrp*, cyan), and myeloid progenitors (*spi1b*, magenta) in bud and early segmentation staged zebrafish (*D. rerio*) (A) and (C) and killifish (*N. furzeri*) (B) and (D) embryos. Dashed boxes indicate zoomed-in regions. Filled magenta arrowheads mark the *spi1b*^+^ ALPM myeloid progenitor domain in *D. rerio*; open arrowheads mark the corresponding region in *N. furzeri*. (**E**) Still image from *in toto* light-sheet imaging of a 10-somite (120 hpf) *N. furzeri* embryo expressing *etsrp*:*mScarlet* (cyan) and ubiquitous *actb*:*membrane-mNeonGreen* (orange). Dashed boxes indicate regions shown in (F) and (G). (**F** and **G**) Zoomed-in views of the ALPM (F) and the yolk (G) showing *etsrp*^+^ vascular cells (cyan arrowheads) and *etsrp*^-^ myeloid cells (orange arrowheads). (**H** and **I**) *In situ* HCR marks *nkx2.7*, *etsrp*, and *spi1b* in *N. furzeri* embryos treated from bud to 10-somites with DMSO control (H, n = 20/20) or 250 nM retinoic acid (I, n = 23/28), imaged at 10-somites. Zoomed-in ALPM regions shown in the panels to the right. Filled white arrowheads mark *etsrp*^+^ LPM cells; open white arrowheads mark the loss of the *etsrp^+^* ALPM domain. ss, somite stage; ALPM, anterior lateral plate mesoderm; RA, retinoic acid. Scale bars: 100 µm (B), (D), (E), (H), and (I), 50 µm (A), (C), and zoomed views.

To follow ALPM dynamics *in vivo*, we imaged *N. furzeri* from the bud to the 10-somite stage using a ubiquitous membrane marker (*40*) and a zebrafish *etsrp* reporter that preferentially labels vascular cells (*41*). As observed by *in situ* HCR, *etsrp*^+^ LPM cells formed bilateral stripes during segmentation (Fig. 4, E and F, and movie S5). Rather than remaining as a compact ALPM domain, a subset of these cells delaminated from the ALPM as chain-like formations that subsequently organized into the yolk sac vasculature (Fig. 4, E to G, and movie S5). Cells exhibiting amoeboid motility were rarely observed emerging from the ALPM (movie S5). Instead, the early myeloid population (*etsrp*^-^) was highly migratory and dispersed throughout the yolk (Fig. 4, E and G, and movie S5). Thus, unlike zebrafish, the *N. furzeri* ALPM is not a major source of myeloid cells.

To directly test whether the ALPM is dispensable for myeloid development in *N. furzeri*, we treated embryos with exogenous retinoic acid during the window of ALPM specification, a perturbation previously shown to disrupt ALPM-derived vascular and myeloid progenitor formation in zebrafish (*42*, *43*). Retinoic acid strongly reduced or abolished the ALPM *etsrp*^+^ vascular progenitor domain, while posterior LPM *etsrp* expression was maintained (Fig. 4, H and I). Despite this disruption, *spi1b*^+^ myeloid cells persisted on the yolk (Fig. 4, H and I). Together, these results demonstrate that primitive myelopoiesis is uncoupled from the ALPM in killifish, indicating that the conserved myeloid GRN can be deployed outside the ventral mesoderm in a reorganized morphogenetic environment.

## Discussion

Gene regulatory networks are typically understood as the primary drivers of development—transcriptional hierarchies whose conserved logic is thought to explain why cell types arise in stereotyped sequences (*1*). Less appreciated is that morphogenetic context may itself constrain when and where these networks are deployed (*44*). Here we show that primitive myelopoiesis has been uncoupled from the ventral mesoderm, the tissue from which it normally arises in vertebrates. Instead, neutrophils first emerge in a salt-and-pepper pattern around the incipient aggregate, spatially separated from the forming mesendoderm. This differentiation occurs while blastomeres remain dispersed across the yolk surface, before gastrulation establishes coherent germ layer tissues. As in other vertebrates, these early neutrophils depend on BMP signaling (*11, 13*). Unlike in other vertebrates, these cells persist when mesendoderm is disrupted. Although neutrophil identity and function are largely conserved, the embryonic architecture in which they arise has substantially changed.

The relocation of primitive myelopoiesis to the aggregate periphery in annual killifish illustrates how conserved developmental programs can be redeployed while preserving components of their core regulatory logic. These findings suggest that myeloid specification is not intrinsically linked to a particular tissue of origin but instead depends on the signaling environment in which the program is activated. The successive waves of vertebrate myelopoiesis, each arising from distinct anatomical locations yet retaining conserved regulatory logic, indicate that the myeloid GRN has been repeatedly redeployed during evolution (*45–48*). Moreover, ectopic BMP signaling is sufficient to activate the myeloid GRN in Xenopus animal cap explants, independent of mesoderm (*49*). In annual killifish, the dispersion of deep cells may create a signaling environment at the aggregate periphery where cells outside the organizer remain exposed to BMP signaling while experiencing lower Nodal activity. Geometry-dependent retention of Nodal ligands within the forming aggregate, as recently proposed in zebrafish (*50*), could generate a steep signaling gradient that confines Nodal activity to the high-density organizer while permitting BMP-dependent myeloid specification at the periphery.

By uncoupling primitive myelopoiesis from the ventral mesoderm, annual killifish provide a new perspective on the proposed common origin of myeloid and vascular lineages. In zebrafish, myeloid and vascular progenitors arise in close proximity within the ALPM, as part of a shared regulatory program involving Etsrp, Tal1, and Alk8-mediated BMP signaling (*11, 13*). This overlap has long complicated efforts to determine whether myeloid and vascular progenitors arise from a shared bipotent progenitor or are independently specified within a common signaling environment (*13*, *51–53*). Killifish serve as a natural experiment that disentangles these normally intertwined lineages: vascular progenitors arise from within the ALPM, whereas neutrophils are specified before germ-layer segregation and outside mesodermal tissue. In other vertebrates, the coordinated emergence of myeloid and vascular progenitors may therefore reflect shared exposure to BMP signaling in the ventral mesoderm rather than a shared developmental origin.

Our findings demonstrate that cell type evolution can occur not only through changes to GRNs themselves, but also through changes in the morphogenetic landscapes in which those networks unfold. In a species adapted to ephemeral environments and prolonged diapause, early immune deployment may confer a selective advantage during an otherwise vulnerable period of developmental stasis. At the same time, the unusual dispersed morphogenesis of annual killifish may have secondarily relocated the myeloid program by altering the organization of BMP and Nodal signaling. More broadly, these results suggest that a cell type can retain its identity even as the developmental lineage that deploys it is evolutionarily reorganized. After all, nature selects for developmental outcomes, not the specific morphogenetic route used to achieve them.

## Acknowledgments

We thank the Shendure lab for providing PIPseq equipment, Saulius Sumanas for sharing the zebrafish *etsrp* promoter plasmid, David Kimelman for sharing the NLS-kikume plasmid, and Elliot Hagedorn for helpful discussions. We thank the Abitua lab members for helpful discussions and critical reading of the manuscript as well as Jessi Gauvin and Laura Stump for animal husbandry.

## Funding

The Searle Scholars Program (PBA)

The Royalty Research Fund (PBA)

National Institutes of Health Genome Training Grant (SMS) (T32HG000035)

The Bonita and David Brewer Fellowship (SMS)

The Shurl and Kay Curci Foundation Ph.D. Fellowship (SMS)

Damon Runyon Postdoctoral Research Fellowship (NCL)

National Science Foundation Graduate Research Fellowship Program (CLTB)

Fluent Biosciences for an in-kind contribution of the PIP-seq scRNA-seq kit and sequencing.

## Author contributions

SMS and PBA conceived the study. SMS designed and performed all experiments. SMS, NCL, and PBA interpreted the results. SMS, NCL, SVH, and CLTB developed methodology. NCL and SMS analyzed the light-sheet data and generated figures. SVH developed the SKiRA desktop app. SMS and PBA wrote the manuscript and it was reviewed by all authors.

## Data, code, and materials availability

Sequencing data have been deposited to the NCBI Gene Expression Omnibus (GSE333732). All code used for analysis of lightsheet, *in situ* HCR, scRNA-seq, and the development of SKiRA has been deposited to GitHub (https://github.com/nlammers371/killi-immune-paper, https://github.com/sydsatsci/HCR_furzeri, https://github.com/sydsatsci/scRNAseq_furzeri, https://github.com/galaxygoldfish/SKiRA). SKiRA can also be downloaded from https://www.abitua.org. Plasmids and transgenic lines are available upon request.

## Supplementary Materials

Materials and Methods

Figs. S1 to S6

Tables S1 to S8

References (*1–61*)

Movies S1 to S5

