## Supplementary Materials for "Early immune cell development precedes gastrulation in annual killifish"

**The PDF file includes:**

Materials and Methods
Figs. S1 to S6
Captions for Tables S1 to S8
Captions for Movies S1 to S5
References

**Other Supplementary Materials for this manuscript include the following:**

Tables S1 to S8
Movies S1 to S5

### Materials and Methods

Animal husbandry and staging

An inbred strain (GRZ) of *Nothobranchius furzeri*, originating from Gonarezhou National Park, Zimbabwe, and an inbred strain (AB) of *Danio rerio* were used in experiments. Both species were maintained at 28°C under 15:9 light:dark cycles with 1-hour light ramp-up and ramp-down periods. Zebrafish staging followed (*54*). Breeding and maintenance of fish followed the Institutional Animal Care and Use Committee (IACUC) protocol, approved by the University of Washington Office of Animal Welfare. Killifish husbandry was performed as previously described (*6*), and zebrafish husbandry followed standard laboratory procedures.

Embryo dissociation and single cell RNA-sequencing

Killifish were crossed for 2 hours. Using the midpoint as 0 hours post fertilization, clutches from 52, 72, 96, and 115 hpf consisting of 132, 90, 80, and 35 embryos, respectively, were visually confirmed to be at dispersed, incipient aggregate, bud, and 8-somite stages. Embryos were manually dechorionated with forceps in ice-cold DMEM/F12 (Gibco, 21-041-025) and transferred to low-bind 1.5 mL Eppendorf tubes on ice. Cells were spun down at 500 x g for 3 min at 4°C. Cells were dissociated in Accutase (Innovative Cell Technologies, AT104) at 37°C for 7-20 min depending on embryo stage (older embryos required longer dissociation). Percent viability was determined by counting $DAPI^+$ (Thermo Fisher Scientific, D1306), Calcein-$AM^+$ (Invitrogen, C1430) live cells and $DAPI^+$, Calcein-$AM^-$ dead cells on a disposable hemocytometer (Bulldog Bio, DHC-N420). 40,000 cells were loaded onto the V4 T20 PIP-seq kit (Fluent BioSciences, now the Illumina Single Cell 3'RNA Prep Kit), except for the 115 hpf, 8-somite stage, which consisted of a 1:1 mix of killifish and stage-matched zebrafish cells. The mixed-species barnyard sample was used to initially assess the efficacy of the PIP-seq kit which, at that time, had not yet been used in teleosts.

scRNA-seq QC and analysis

cDNA fragment analysis of scRNA-seq libraries was performed using the 4200 TapeStation System. Libraries were sequenced with a NextSeq 2000 P3 100 cycle kit for the 52 hpf, 72 hpf, and 96 hpf timepoints, or with a NovaSeq 6000 S4 300 cycle kit for the 115 hpf timepoint using the following configuration: read 1: 54, i7 index: 8, i5 index: 8, read 2: 68. Reads were aligned to the *N. furzeri* (Nfu_20140520) or the *D. rerio* (GRCz11.108) genomes, and gene-count matrices were generated using PIPseeker software (v1.1.7, Fluent BioSciences). For the 115 hpf timepoint only, cells with >90% of reads mapping to the killifish genome were labeled killifish cells and computationally isolated from the killifish-zebrafish mix. Doublets were removed using Scrublet (v0.2.3). Cells with >12% mitochondria reads and >5000 reads/cell were removed. Downstream analysis was performed with Seurat (v5.4.0). On average, the number of reads per cell and genes per cell were as follows: 52 hpf, 11,366 reads, 3,182 genes; 72 hpf, 12,738 reads, 3,246 genes; 96 hpf, 10,387 reads, 2,886 genes; 115 hpf, 7,876 reads, 2,524 genes. The R notebooks are available at https://github.com/sydsatsci/scRNAseq_furzeri.

Small molecule perturbations

To inhibit Activin/Nodal receptors (ALK4, ALK5, and ALK7), *N. furzeri* embryos were treated with 1 µM A-83-01 (Sigma, SML0788). To inhibit BMP receptors (ALK1, ALK2, ALK3, and ALK6), embryos were treated with 40 µM of LDN-193189. Exogenous retinoic acid (RA) was applied at 250 nM from 96 to 120 hpf. A-83-01 and LDN-193189 treatments were applied from 48 to 72 hpf, washed out, and then at 96 hpf (A-83-01 and LDN-193189) or at 120 hpf (RA) the embryos were either imaged on the light sheet or fixed. For fixation, embryos were incubated in 4% paraformaldehyde (Fisher Scientific, 50-980-495) for 1 hour at room temperature and then overnight at 4°C. Embryos were rinsed in 0.05% Tween-20 (Sigma, P1379) in PBS (Fisher Scientific, AM9625) before a methanol dehydration series for HCR/IHC.

*In situ* HCR and IHC

Hybridization chain reaction (HCR) probes were designed using the *in situ* probe generator Jupyter notebook (https://github.com/rwnull/insitu_probe_generator) (*55*) and ordered as oPools from IDT (table S8). HCR reagents were purchased from Molecular Instruments and were used according to the manufacturer's protocol, with modifications from the supplementary information for zebrafish whole mount HCR (*56*). In brief, 10-20 embryos for each sample were rehydrated from 100% methanol to 100% 0.05% Tween-20 (Sigma, P1379) in PBS (Fisher Scientific, AM9625) through a graded series of washes. For each sample, 1 pmol of each probe was added to 250 µL of hybridization buffer (Molecular Instruments), and embryos were incubated at 37°C overnight. Embryos were then washed with wash buffer, and 2.5 µL of each snap-cooled hairpin amplifier (B1-488, B3-546, B5-647) was added to 250 µL of amplification buffer and incubated at room temperature overnight. For IHC, phospho-SMAD1/5/9 rabbit mAb (Cell Signaling Technology, 13820T) and secondary anti-rabbit Alexa Fluor 488 antibody (Thermo Fisher, A11304) were used at 1:1000 dilution with 0.05% Tween-20 in PBS. Embryos were stained with DAPI (Thermo Fisher Scientific, D1306) for 1 hour, as recommended by the manufacturer.

Stable transgenesis

Four stable lines were generated using the Tol2 system or CRISPR-Cas9, as previously described (*57*, *58*). The nuclear fluorescent marker Tol2(*actb*:*NLS-mScarlet*) was subcloned from a previous study (*59*) into the Tol2 vector, driven by the 1316 bp β-actin promoter. The vascular progenitor reporter Tol2(*etsrp*:*mScarlet*) was generating using a plasmid gifted by Saulius Sumanas from a previous study (*41*), which contains a 2.3 kb region of the zebrafish *etsrp* promoter. The promoter was subcloned into the Tol2 vector driving mScarlet. The membrane fluorescent marker Tol2(*actb*:*membrane-mNeonGreen*) was obtained from a previous study (*40*). For the CRISPR-KI line (*eGFP-P2A-lcp1*), eGFP and a self-cleaving P2A site were amplified with 35 bp of homology to the N-terminus of Lcp1, serving as the homology donor template (HDT) for homology-directed repair. The injection mix consisted of 10 ng/µL HDT, 50 ng/µL RNP (AltR CRISPR-Cas9 + sgRNA [crRNA+tracrRNA]; IDT, 1081058; IDT, 1072532), 1 µM HDR enhancer (IDT, 10007910), and 1X phenol red (Sigma, P0290-100ML). Genomic DNA was extracted using the HotSHOT method, as previously described (*60*), and subsequent genotyping via PCR and Sanger sequencing confirmed a complete in-frame insertion of eGFP-P2A at the N-terminus of Lcp1. Oligonucleotide sequences are provided in table S8. All plasmids and transgenic lines are available upon request.

Imaging

For confocal imaging, embryos were mounted in 1.2% low-melting-point agarose (Fisher Bioreagents, BP1360) in PBS (Fisher Scientific, AM9625) in 60-mm Petri dishes (Millipore Sigma, P5481). Confocal images were acquired on a Zeiss LSM 900 upright microscope using

W Plan-Apochromat 10X or 20X objectives. Confocal stacks were typically 150-300 µm thick. For live imaging, embryos were mounted in glass capillaries (Zeiss, 402100-9330-000) containing 1.2% low-melting-point agarose in DI water and imaged on a Zeiss Lightsheet 7. The light sheet chamber was filled with filtered aquarium water containing 100 units/mL of penicillin-streptomycin (Fisher Scientific, 15-140-122) to prevent bacterial growth. Images were captured at intervals of either 60, 90, or 120 seconds from a single- or dual-sided view. Z-stacks were 650 µm thick with a 3 µm step size to provide sufficient overlap for subsequent computational dual-side stitching.

Cell tracking pipeline

Preprocessing and segmentation of light-sheet movies were performed using custom image analysis software implemented in Python. Raw datasets were exported in Zarr format. Automated dual-view stitching was performed using cross-correlation of voxels in overlapping regions from each side. Nuclear segmentation was performed using a method based on Laplacian-of-Gaussian filtering. eGFP-*lcp1* fluorescence intensity was obtained by calculating the average fluorescence within each nuclear mask. Live-cell tracking was performed using the Ultrack software package (*61*). For the depicted lineage tree, manual track correction and stitching were performed using custom curation software. Deep-cell density on the embryo surface was estimated using the binned cell counts on the surface of a sphere fit to the embryo surface. Counts were smoothed using a Gaussian kernel with sigma = 50 µm. Cells were classified as *lcp1*$^+$ if their average per-frame eGFP-*lcp1* intensity exceeded the 99th percentile of stage-matched deep cells. For the wound assay, net migration velocity of *lcp1*$^+$ cells toward and away from the wound site was calculated using manually annotated, time-resolved wound locations. Cell velocities were denoised using a Savitzky–Golay filter with a 5-minute window length and were averaged over spatial regions of $\pi/10$ radians (18 degrees). The pipeline is available at https://github.com/nlammers371/killi-immune-paper.

*In situ* HCR + IHC quantification

Maximum-intensity projected CZI images were analyzed in Python using NumPy (v1.26.4), pandas (v2.3.3), Matplotlib (v3.10.8), SciPy (v1.15.2), scikit-image (v0.25.2), and AICSImage (v4.10.0). Multichannel images were loaded in CYX format, and CZI voxel metadata were used to convert pixel distances to micrometers. Nuclei were segmented from the DAPI channel by Gaussian smoothing (sigma = 1), Otsu thresholding with objective-specific scaling, small-object removal, hole filling, and seeded watershed segmentation on the Euclidean distance transform. Segmented objects were further filtered by area, solidity, and eccentricity to exclude yolk syncytial nuclei and enveloping layer nuclei and retain deep cells. Mean intensity was measured in each channel for each nuclear mask. Channel background was estimated from the dimmest 20% of nuclei, subtracted from mean nuclear intensities, and negative values were clipped to zero. For radial profiles, a patterning center was manually defined from Nodal expression, and nuclear centroid distances from this center were calculated. In whole embryo 10X images, the embryo boundary was estimated by identifying the farthest nucleus within angular sectors and fitting these points with a smooth closed spline. Per-nucleus signal was plotted as normalized mean nuclear intensity as a function of centroid distance from the patterning center. Radial trends were summarized with rolling 10$^{th}$-90$^{th}$ percentile envelopes. The Jupyter notebook is available at https://github.com/sydsatsci/HCR_furzeri.

Light-sheet quantification

For inhibitor-treated embryos, *lcp1*$^+$ neutrophils were manually counted from *in toto* light-sheet stills of Tol2(*actb*:*NLS-mScarlet*); *eGFP-lcp1* embryos at bud stage (96 hpf). For each embryo, the total number of *lcp1*$^+$ cells was calculated by summing all detected *lcp1*$^+$ cells with a discernible *NLS-mScarlet*$^+$ nucleus across the imaged embryo. Statistical comparisons were performed in R using two-sided Wilcoxon rank-sum tests for planned vehicle-control comparisons. P values were adjusted for multiple comparisons using the Holm method.

Stab wound assay

Using a micromanipulator, a glass capillary needle (Harvard Apparatus, GC100F-10) commonly used for one-cell injections was used to inflict a single stab wound in the posterior region of tailbud stage embryos aiming for Kupffer's vesicle or the yolk. Embryos were transferred to the light-sheet microscope and imaged as described above within 20 minutes of wounding or embryos were fixed 1.5 hours post-wounding for subsequent *in situ* HCR and imaging.

Single-cell Killifish RNA-sequencing Atlas (SKiRA) desktop application

A custom desktop application, SKiRA, was developed to enable interactive exploration of the *N. furzeri* single-cell RNA sequencing embryogenesis atlas, which includes the four developmental timepoints and a merged dataset combining all timepoints. This application allows users to generate UMAPs of gene expression or cell types across each timepoint. Plots can be generated on demand, customized by color, DPI, and labeling, and exported locally in PNG, SVG, or PDF formats. Morphological context is also provided through DAPI-stained images and schematic references of *N. furzeri* embryos at each timepoint. SKiRA was developed using the cross-platform Jetpack Compose Desktop framework (v1.9.2) built on Kotlin (v2.1.20) and compiled for Windows 10 and later and macOS 11 Big Sur and later. Backend data processing was implemented using custom R scripts (v4.5.2) and Seurat (v4.4.0), which interact with the application's user interface to render the relevant plots. SKiRA, including the source code, is open-source and freely available under the Apache 2.0 license. The user installation guide and application are available on GitHub (https://github.com/galaxygoldfish/SKiRA) and on our lab website (https://www.abitua.org/).

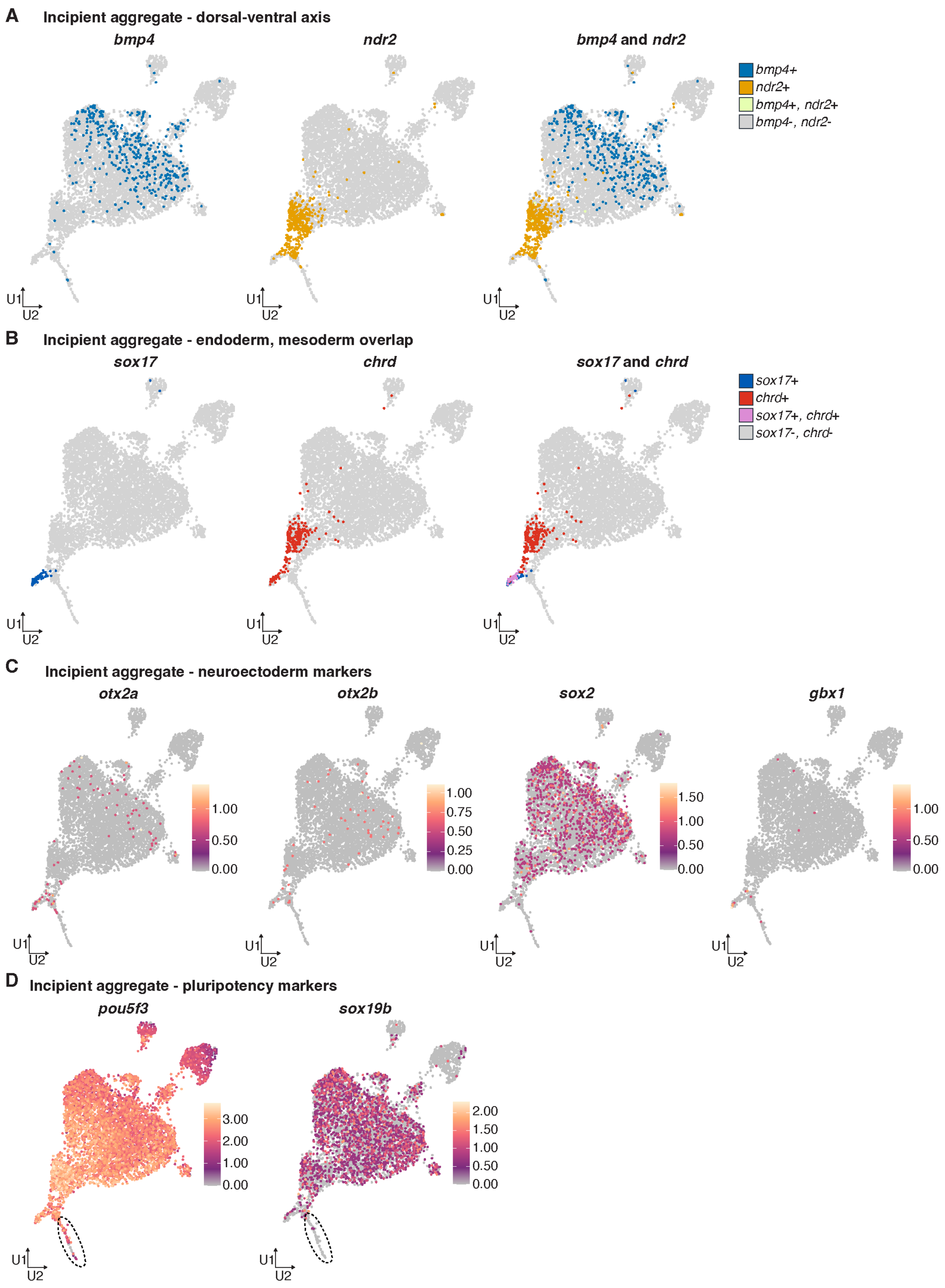


**Fig. S1. Incipient aggregate expression patterns.**
(**A** to **D**) UMAPs of cells from the incipient aggregate stage (72 hpf) showing expression of *bmp4* (blue), *ndr2* (orange), and coexpression (green) in (A), *sox17* (blue), *chrd* (red), and

coexpression (pink) in (B), neuroectoderm markers *otx2a*, *otx2b*, *sox2*, and *gbx1* in (C), and pluripotency markers *pou5f3* and *sox19b* in (D). The dotted black line encircles the pre-gastrula myeloid lineage in (D).

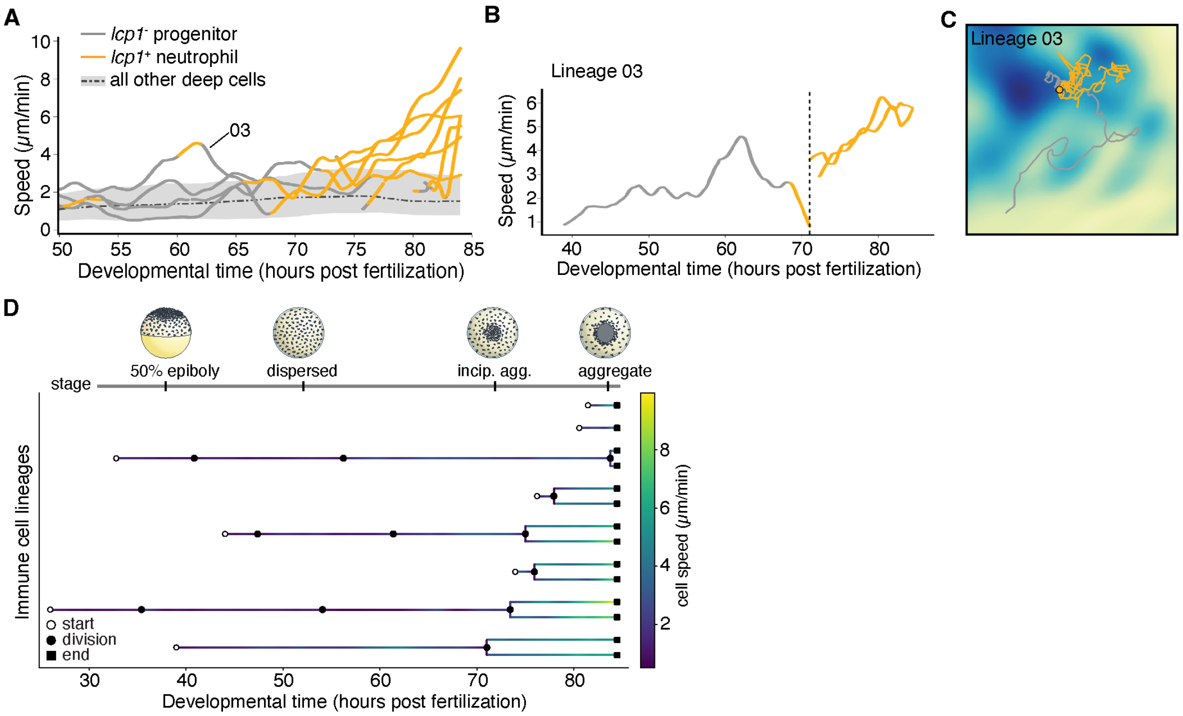


**Fig. S2. Pre-gastrula neutrophil speed over developmental time.**
(**A**) Average speed (μm/min) for all deep cells (dotted line) with minimum and maximum speeds (grey shading) over development. Individual myeloid lineage speeds shown before (grey line) and after (orange line) GFP-*lcp1* expression exceeds background. (**B**) The speed of an individual myeloid lineage is shown over development, including the division event (dotted black line) that produces a sibling myeloid cell pair. (**C**) The location of the myeloid lineage in (B) is shown relative to the local deep cell density (low density – yellow, high density – dark blue). (**D**) Immune cell lineages colored by speed (μm/min), lineage start (open circle), cell division event (closed circle), lineage end (closed square).

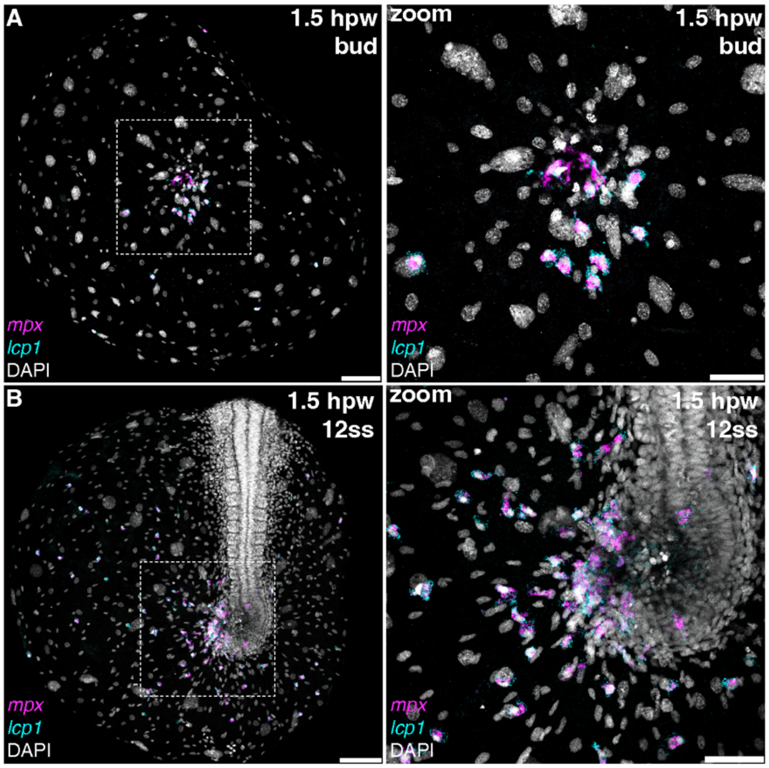


**Fig. S3. Neutrophils coexpressing *mpx* and *lcp1* respond to wound sites.**

(**A** to **C**) *In situ* HCR marks mature neutrophils (*mpx*, magenta; *lcp1*, cyan) localized to a wound site 1.5 hpw. Wounds were inflicted at the ventral yolk (A) or dorsal tailbud (B) at either the bud (A) or the 12-somite stage (12ss) (B). Dashed white boxes indicate zoomed-in views. hpw, hours post wounding; ss, somite stage. Scale bars 100 µm (A) and (B), 50 µm (zoom).

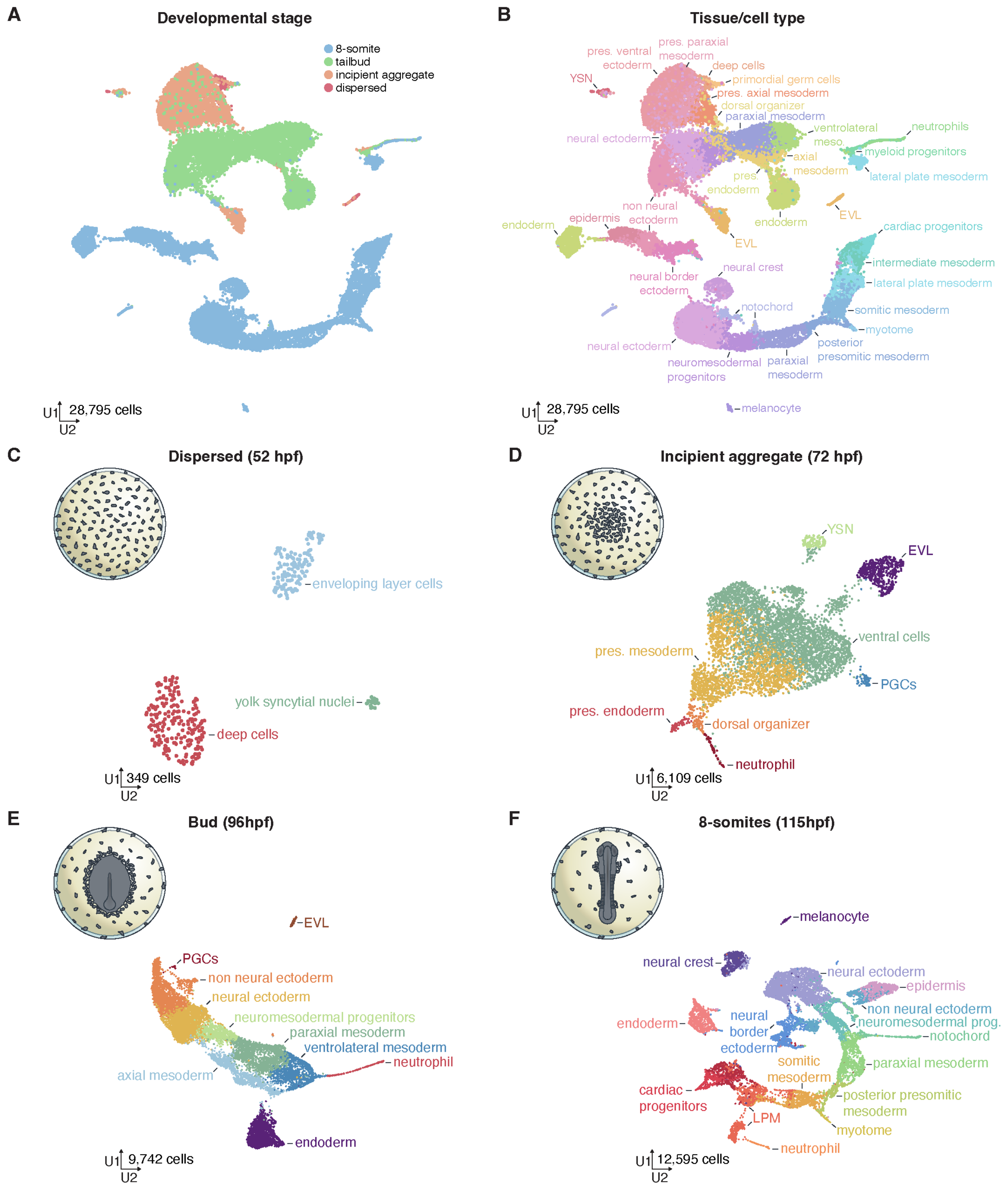


**Fig. S4.** ***N. furzeri*** **embryogenesis scRNA-seq atlas.**
(**A** to **F**) UMAPs show cells from all timepoints colored by timepoint (A), tissue/cell type (B), or individual timepoints from the dispersed (C), incipient aggregate (D), bud (E), or 8-somite (F) stage. Pres. (presumptive), meso. (mesoderm), prog. (progenitor), EVL (enveloping layer cell), YSN (yolk syncytial nuclei), PGCs (primordial germ cells), LPM (lateral plate mesoderm).

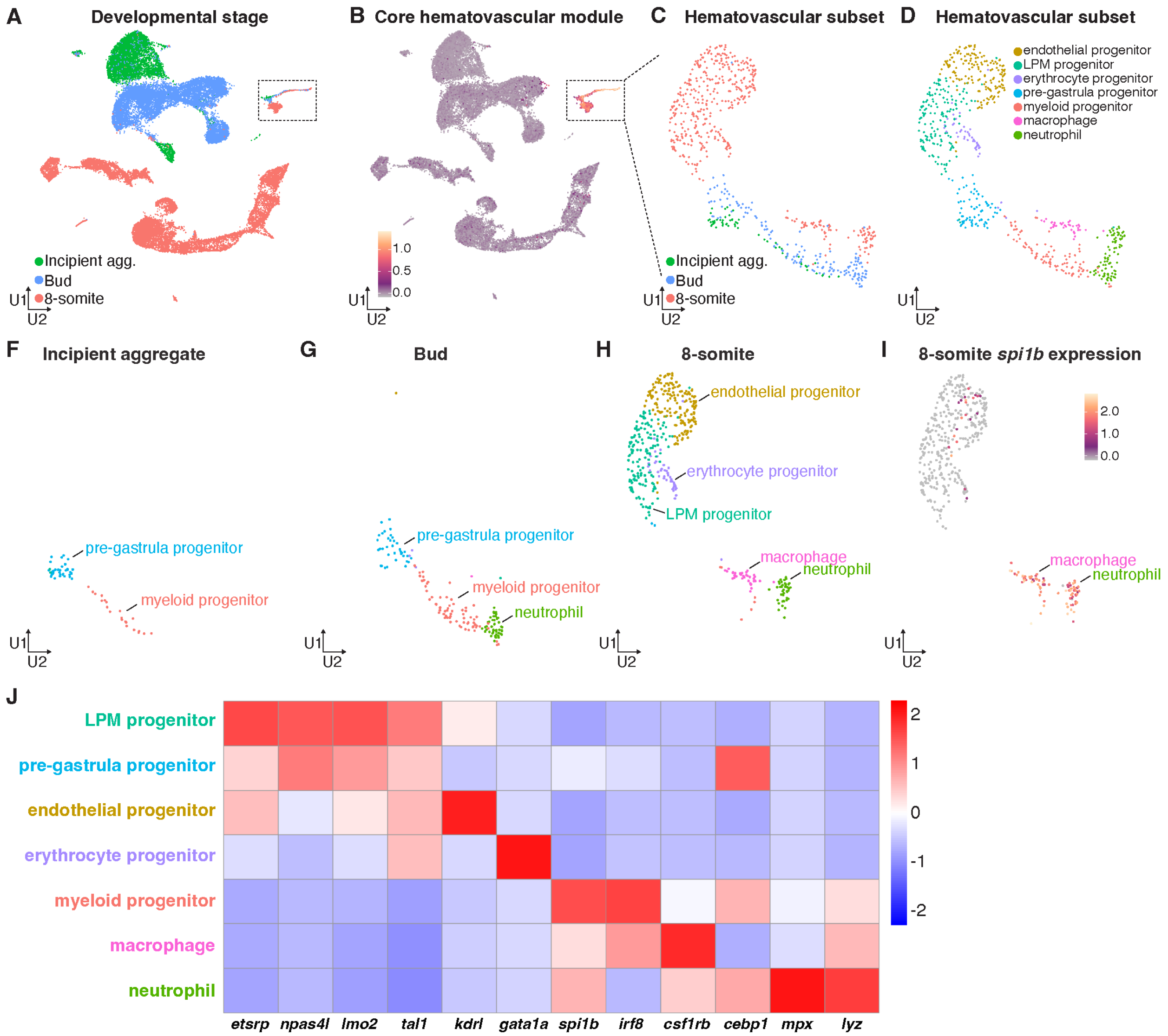


**Fig. S5. Hematovascular scRNA-seq cell types and expression patterns.**

(**A** and **B**) UMAPs show cells from incipient aggregate (72 hpf), bud (96 hpf), and 8-somite (115 hpf) stage *N. furzeri* embryos colored by developmental stage in (A) or the expression of core hematovascular markers in (B) listed in (J). Dotted black rectangle shows the subset of cells in (C). (**C** to **I**) UMAPs show a subset of hematovascular cells from (B) colored by developmental stage (C), cell type (D) or cell type and separated by developmental stage (F, incipient aggregate; G, bud; H, 8-somite). Expression of *spi1b* marks macrophages and neutrophils at the 8-somite stage in (I). (**J**) Gene expression z-score by gene (column) for each cell type (row) for the indicated marker genes.

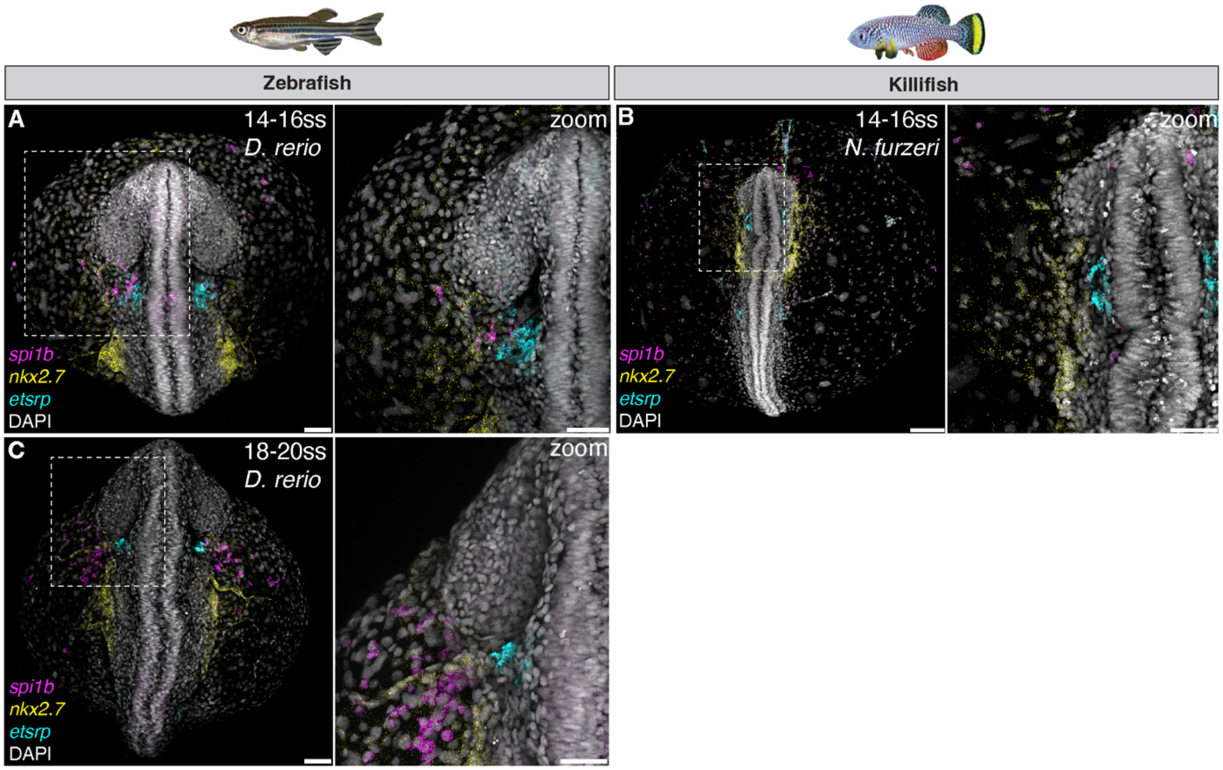


**Fig. S6. The ALPM in *N. furzeri* and *D. rerio*.**

(**A** to **C**) HCR *in situs* mark the ALPM (*nkx2.7*, yellow), vascular progenitors (*etsrp*, cyan), and myeloid progenitors (*spi1b*, magenta) in zebrafish (*D. rerio*) in (A) and (C) and killifish (*N. furzeri*) embryos in (B) at late segmentation stages. Dashed white boxes indicate regions shown in zoomed-in views of the ALPM in the adjacent panels. Scale bars 50 µm (A) and (C) and zoomed views, 100 µm (B).

**Table S1. Differentially expressed genes for incipient aggregate stage (72 hpf) scRNA-seq clusters used for the identification of cell type enriched genes.**

All genes listed are differentially expressed in the cell type listed in the "cluster" column as compared to all other cells in the dataset. "p_val" is the p-value, "avg_log2FC" is the average log 2-fold change (positive – upregulated, negative – downregulated), "pct.1" and "pct.2" represent the percent of cells within the cell type listed in the "cluster" column expressing the given gene in the "gene" column.

**Table S2. Differentially expressed genes for bud stage (96 hpf) scRNA-seq clusters used for the identification of cell type enriched genes.**
Columns as defined in table S1 caption.

**Table S3. Differentially expressed genes for 8-somite stage (115 hpf) scRNA-seq clusters used for the identification of cell type enriched genes.**
Columns as defined in table S1 caption.

**Table S4. Differentially expressed genes for dispersed stage (52 hpf) scRNA-seq clusters used for the identification of cell type enriched genes.**
Columns as defined in table S1 caption.

**Table S5. Differentially expressed genes for scRNA-seq clusters from all stages (52, 72, 96, 115 hpf) used for the identification of cell type enriched genes.**
Columns as defined in table S1 caption.

**Table S6. Canonical marker genes used to identify the hematovascular lateral plate mesoderm subset in fig. S5.**

**Table S7. Differentially expressed genes for hematovascular lateral plate mesoderm subset scRNA-seq clusters (72, 96, 115 hpf) used for the identification of cell type enriched genes.**
Columns as defined in table S1 caption.

**Table S8. Oligonucleotides used in this study.**
Includes primers, gRNAs, and HCR probes.

**Movie S1.**
This movie shows a transgenic *N. furzeri* embryo expressing *eGFP-lcp1* myeloid marker (orange) and *actb*:*NLS-mScarlet* nuclear marker (white) from the start of epiboly (26 hpf) to the late aggregate stage (84.4 hpf). The time-lapse first shows the animal side of the embryo, then the vegetal side and is approximately 60 hours long with an interval of 90 seconds. (MPEG; 281.1 MB).

**Movie S2.**
This movie shows a 2D Mercator projection of the *N. furzeri* embryo in movie S1. Tracks show *lcp1*$^-$ progenitors (black track, black circle), transitioning (orange star) to *lcp1*$^+$ neutrophils (orange track, orange circle). Nuclear density is displayed topographically where yellow is low density and dark blue is high density (nuclei/mm$^2$). (MPEG; 10.2 MB).

**Movie S3.**
This movie shows a transgenic *N. furzeri* embryo expressing *eGFP-lcp1* myeloid marker (orange) and *actb*:*NLS-mScarlet* nuclear marker (white) at bud stage (96 hpf) zoomed into the yolk adjacent to the developing embryo. The time-lapse is approximately 18 hours long with an interval of 60 seconds. (QuickTime; 102.5 MB).

**Movie S4.**
This movie shows a transgenic *N. furzeri* embryo expressing *eGFP-lcp1* myeloid marker (orange) and *actb*:*NLS-mScarlet* nuclear marker (white) at bud stage (96 hpf) with a stab wound inflicted near Kupffer's vesicle. Neutrophil tracks toward (red) and away from (blue) the wound are shown. The time-lapse is approximately 12 hours long with an interval of 60 seconds. (MPEG; 6.8 MB).

**Movie S5.**
This movie shows a transgenic *N. furzeri* embryo expressing *etsrp*:*mScarlet* (hematovascular progenitor marker; white) and *actb*:*membrane-mNeonGreen* (membrane marker; magenta) from the bud stage to the 10-somite stage. The time-lapse is approximately 24 hours long with an interval of 120 seconds. (QuickTime; 69.3 MB).

56. H. M. T. Choi, C. R. Calvert, N. Husain, D. Huss, J. C. Barsi, B. E. Deverman, R. C. Hunter, M. Kato, S. M. Lee, A. C. T. Abelin, A. Z. Rosenthal, O. S. Akbari, Y. Li, B. A. Hay, P. W. Sternberg, P. H. Patterson, E. H. Davidson, S. K. Mazmanian, D. A. Prober, M. van de Rijn, J. R. Leadbetter, D. K. Newman, C. Readhead, M. E. Bronner, B. Wold, R.

Lansford, T. Sauka-Spengler, S. E. Fraser, N. A. Pierce, Mapping a multiplexed zoo of mRNA expression. *Development* **143**, 3632–3637 (2016).
